# Bacterial vitamin B6 required for post-embryonic development in *C. elegans*

**DOI:** 10.1101/2023.10.18.562920

**Authors:** Min Feng, Baizhen Gao, Daniela Ruiz, L. Rene Garcia, Qing Sun

## Abstract

Nutritional intake influences animal growth, reproductive capacity, and survival of animals. Under nutrition deficiency, animal developmental arrest occurs as an adaptive strategy to survive. However, the nutritional basis and the underlying nutrient sensing mechanism essential for initiating animal regrowth after developmental arrest remain to be explored. In *Caenorhabditis elegans*, larvae undergo early developmental arrest are stress resistant, and they require certain nutrients to initiate postembryonic development. Here, we investigated the developmental arrest in *C. elegans* feeding on *Lactobacillus plantarum*, and the rescue of the diapause state with trace supplementation of *Escherichia coli*. We performed a genome-wide screen using 3983 individual gene deletion *E. coli* mutants and identified *E. coli* genes that are indispensable for *C. elegans* larval growth on original not nutritionally sufficient bacteria *L. plantarum*. Among these crucial genes, we confirmed *E. coli pdxH*, and the downstream metabolite pyridoxal 5-P (PLP, Vitamin B6) as essential nutritional factors initiating *C. elegans* postembryonic development. Transcriptome results suggest that bacterial *pdxH* affects host development by coordinating host metabolic processes and PLP binding. Additionally, bacterial PLP may act as a cofactor for host tyrosine aminotransferase, thereby promoting the translocation of daf-16 to nucleus. Altogether, these results highlight the role of microbial metabolite PLP as a crucial micronutrient to recover postembryonic development in *C. elegans*.

## Introduction

A well-balanced diet contains macronutrients such as carbohydrates, fats, and proteins, as well as micronutrients such as vitamins and minerals to sustain basal metabolism and tissue function. Lack of certain nutrients leads to alterations in metabolism and causes severe deficiency in animal growth and reproduction. Therefore, understanding the role of individual nutrients is critical for revealing regulatory mechanisms of animal development.

The nematode *Caenorhabditis elegans* is a good model to study the effects of diet on life history traits such as development, reproduction, and aging^1^. As a bacterivore, *C. elegans* consumes a wide range of bacteria, including bacteria found in human microbiota such as *E. coli*. Multiple genetic engineering techniques have been developed to modify the genome of both *C. elegans* and its diet bacteria, making this model suitable for investigating the effects of bacteria-derived nutrients on host development^2^. Nutrient availability is a determinant of larval development in *C. elegans*. To survive unfavorable nutritional environments, the larvae can alter their developmental progression^3^. For example, in the absence of nutrients, newly hatched L1 larvae immediately arrest their postembryonic development^4^. Therefore, the *C. elegans* developmental arrest model provides an opportunity to study how the animal adapt to fluctuations in nutrient availability.

Animals obtain essential nutrients not only from food intake but also from the synthesis of co-existing symbiosis microbiota. Previous studies emphasized the importance of metabolites derived from bacteria on larvae development in *C. elegans*. For example, vitamin B12, a metabolite of *Comamonas aquatica* D1877, accelerates host development via the methionine/S-Adenosylmethionine (SAM) cycle^5^. In addition, bacteria-secreted enterobactin facilitates mitochondrial iron uptake and promotes worm development by binding to the ATP synthase^6^. Additionally, vitamin B2 provided by bacteria regulates worm protease activities and development^7^. However, the effects of microbial metabolites/nutrients on the initiation of postembryonic development in animals are missing, which is essential for understanding both the nutritional requirements and the intricate mechanism involved in animal growth.

In this study, an interspecies systems biology approach with *C. elegans* and its microbial diets, *E. coli* and *Lactobacillus plantarum*, were used to identify bacterial metabolites that affect the animal’s early-stage development and gene expression. We observed that hatched L1 *C. elegans* developmentally arrest when feeding on *L. plantarum* diet; however transient feeding with trace *E. coli* can promote larval development and further feeding on *L. plantarum*, which was originally not nutritionally sufficient. Through a systematic screen on ~4000 single gene deletion strains, we identified 29 *E. coli* genes that are essential from *E. coli* to promote *C. elegans* L1 growth on a *L. plantarum* diet. The *E. coli pdxH* gene and the downstream metabolites vitamin B6 was further confirmed to coordinate host metabolic processes and PLP-binding activity thereby contribute to the host development.

## Results

### *L. plantarum* induces early developmental arrest in *C. elegans*

The nutritional composition of bacterial food can induce larval developmental delay in *C. elegans*. To investigated the worm growth on Gram-positive bacterium *L. plantarum*, a commonly consumed probiotic strain, and compare that with standard laboratory food *E. coli* OP50. Synchronized L1 worms were fed with *L. plantarum* or *E. coli* OP50. We found that ~95% worms fed on OP50 developed to adults within 3 days at 21-22°C, whereas worms fed solely on *L. plantarum* and developmentally arrested at early larval stage (Figure 1A), suggesting that *L. plantarum* may not provide sufficient nutrients for *C. elegans* larvae development. To further investigate food preference of *C. elegans*, two behavior assays to assess food dwelling and food choice assay were performed. Briefly, in the food dwelling assay, we placed worms on a bacterial lawn and observed their location after a 24-hour period. For the food choice assay, we set a plate with OP50 lawn on one side and *L. plantarum* lawn on the opposite side. The percentages of worms found on each bacterial lawn were monitored at specific time intervals to assess their food preference. The results showed worms tend to stay on OP50 as compared to the *L. plantarum* lawn (Figure 1B and C), indicating *L. plantarum* is a less favorable food source. Interestingly, in the food choice assay, after OP50 was consumed at day 4, ~25% worms switched to the *L. plantarum* lawn and consumed it (Figure 1C), suggesting that worms can utilize *L. plantarum* food for growth, if they previously ate OP50.

**Figure 1.**
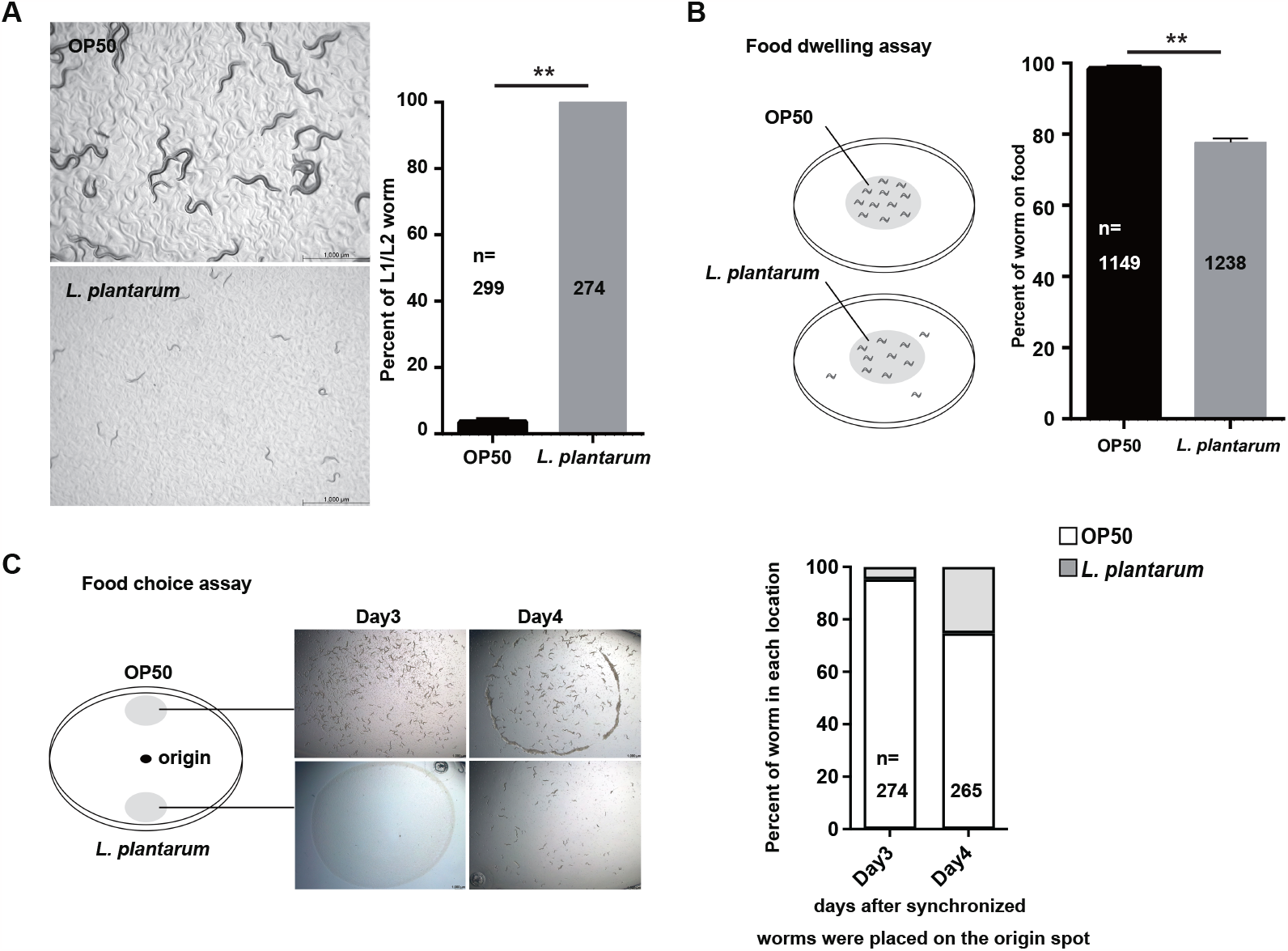

### *L. plantarum* induced development arrest is not due to bacterial avoidance

As *C. elegans* pharyngeal pumping filters bacteria and transports them from the stoma to the isthmus^8^, we then asked if the size of *L. plantarum* (0.9–1.2 µm wide and 3–8 µm long^9^) is too large for L1 worms to feed. In order to see if *L. plantarum* can pass the worms pharynx and reach the gut, we used a green fluorescent dye carboxyfluorescein diacetate succinimidyl ester (CFSE) to stain *L. plantarum* and fed the fluorescent *L. plantarum* to *C. elegans*. After 2 hours of incubation, we found L1 worms fed CFSE stained *L. plantarum* had green fluorescence signal both in the pharynx and the gut (Figure 2A), indicating that the dietary effect from *L. plantarum* on worm development was not due to deficiency of bacteria uptake.

**Figure 2.**
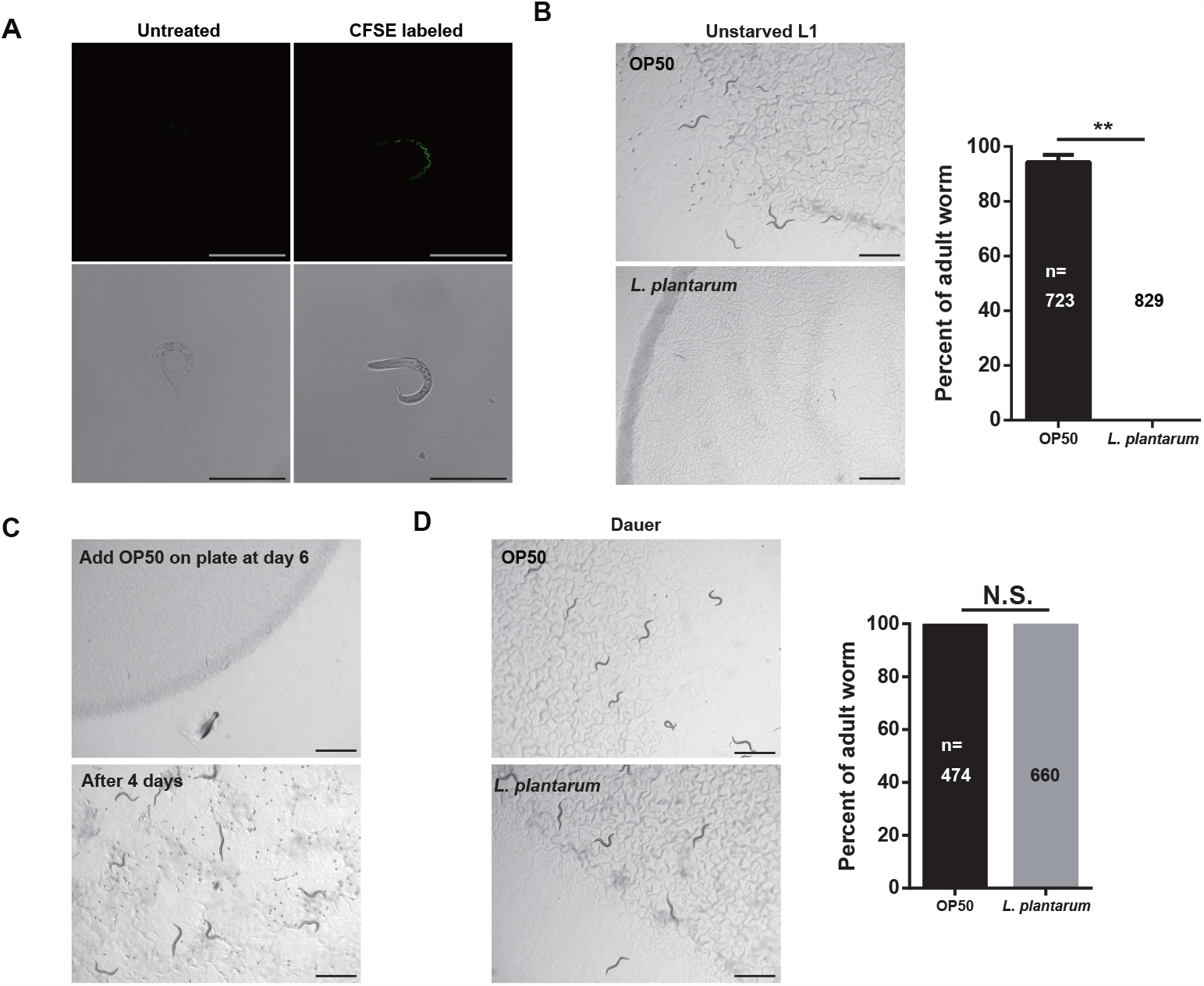

Since starvation causes animal physiological and behavioral changes, we wondered whether these changes are responsible for the developmental arrest in worms fed on *L. plantarum*. To test this hypothesis, newly isolated eggs are placed on the *L. plantarum* lawn to avoid starvation. However, the worms stayed at L1/L2 stage 5 days after been placed on the *L. plantarum* lawn (Figure 2B), suggesting the developmental arrest induced by *L. plantarum* is not because the worm avoided *L. plantarum* after hatching. To explore whether this developmental arrest affect is reversible, OP50 was added to worms exposed to *L. plantarum* for 6 days. After adding OP50, developmental arrested worms recovered to adults with viable progenies within 3 days (Figure 2C). This suggests that the nutrient deficiency of *L. plantarum* induced a reversible developmental arrest to the host, which is a similar protective response to starvation. We next asked if *L. plantarum* is also a nutrient-deficient food source for another typical development arrested stage, dauer larvae. To test this, dauer larvae were isolated and fed either OP50 or *L. plantarum*. We found that dauer worms grew normally into adults when fed *L. plantarum* food (Figure 2D). These findings suggest that *L. plantarum* lacks certain nutrients that are essential for the growth of early-stage worms, but it provides sufficient nutrients for exit from the dauer stage to adulthood.

### A trace amount of *E. coli* can support the development of worm fed on *L. plantarum*

Previous studies have shown that when larvae are provided with inadequate food, larval growth can be restored by supplementing a trace amount (0.2 µl, OD_600_=1, 1.6×10^8^ c.f.u.) of live *E. coli*^6^. To determine whether *E. coli* metabolites can compensate for some *L. plantarum* nutritional deficiency, L1 worms first were fed the same trace amount of live *E. coli* OP50 or BW25113. The optical refractivity of worms fed with 0.2 µl of bacterial culture changed so that they were easier to visualize, but they failed to develop substantial biomass (Figure 3A).

**Figure 3.**
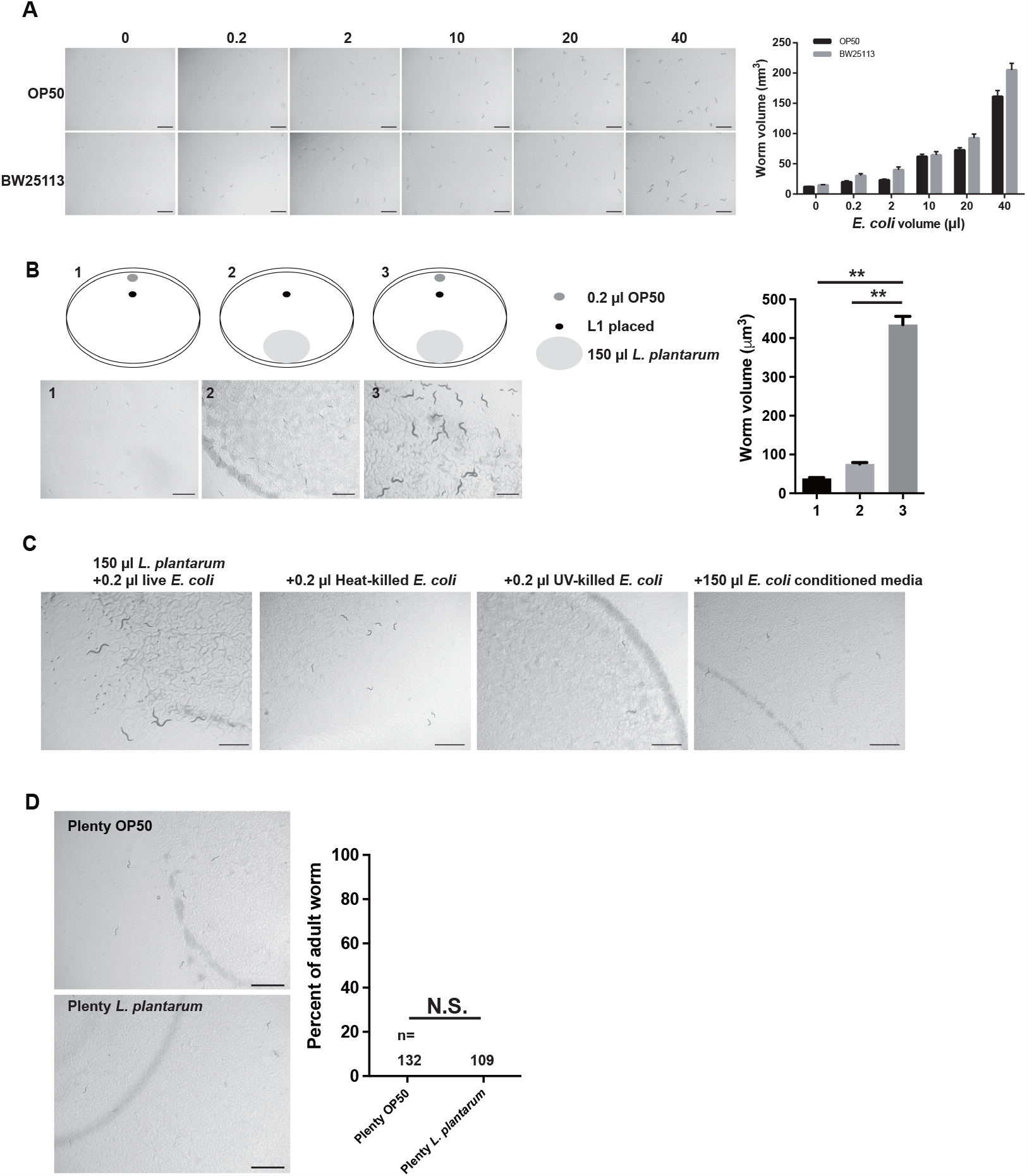

Next, we fed the worms abundant *L. plantarum* together with trace live *E. coli*. We found that larval growth was recovered with the supplemented *E. coli* (Figure 3B), and the progeny from those worms were also able to grow on *L. plantarum*, suggesting that factors from the trace amount of *E. coli* stimulate *C. elegans* surpass L1/L2 stage so they can use *L. plantarum* as nutrients. To characterize further the active compounds from *E. coli*, we supplemented *L. plantarum* with heat-treated, UV-treated *E. coli* or *E. coli*-conditioned media. We observed that while live *E. coli* promoted *C. elegans* growth of worms on *L. plantarum*, none of the above conditions were as effective (Figure 3C). Thus, *E. coli* factors that rendered the *L. plantarum* edible are not secreted, heat nor UV stable. Increasing evidence suggests that various bioactive dietary factors may modify the epigenome and could thus be incorporated into an epigenetic diet^10^. Here we sought to determine whether a diet supplemented with *L. plantarum* could modify the epigenetic patterns of *C. elegans* and render the diet edible for their offspring. To test this, gravid worms were fed trace amounts of *E. coli* with an abundant supply of *L. plantarum*, then bleached to obtain L1 worms, which were subsequently placed on a *L. plantarum* lawn. As a control, L1 worms from mothers fed only *E. coli* were used. After five days on the plate, the worms remained in the early developmental stage, similar to the control group (Figure 3D). These results suggest that potential epigenetic modifications resulting from the *L. plantarum* diet were insufficient to render the diet edible for their offspring.

### A high-throughput screen of *E. coli* single gene deletions identifies the genes required for *C. elegans* to fed and develop on *L. plantarum*

To identify systematically the bacterial components that act as nutrient signals to recover worm development, we performed a genome-wide screen using the Keio collection library. This library contains 3985 single gene knockout *E. coli* strains, with individual genes replaced with a kanamycin cassette in the K12 BW25113 background^11^. To conduct the screen, approximately 100 synchronized L1 worms were placed onto the plates with a trace amount of *E. coli* mutants from the Keio collection, together with *L. plantarum* (Figure 4A). We then monitored the worms’ development for 4-6 days. The wild-type parental strain *E. coli* BW25113 was used as the positive control. In the primary screen, we selected *E. coli* candidates that potentially delayed worm development. We performed a secondary screen with biological triplicates to narrow the candidates to 421 *E. coli* mutants (Figure 4A). After a third screening, we confirmed 29 mutant *E. coli* strains that fail to support worm growth (Figure 4B, Table 1).

**Figure 4.**
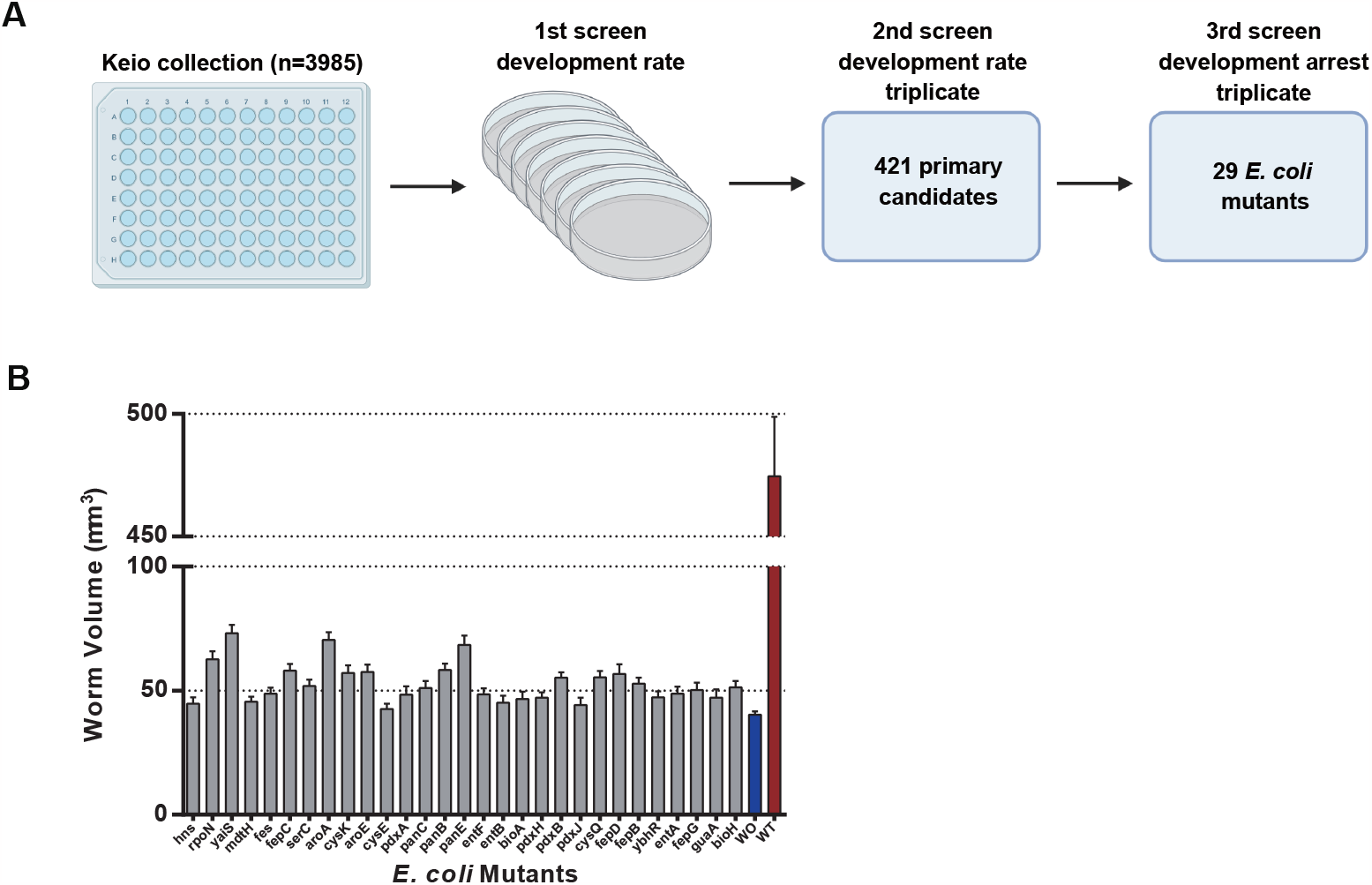

### *E. coli* vitamin B6 is essential for post-embryonic development in *C. elegans*

To classify the function of these 29 genes essential for recovering *C. elegans* early larval stage development, we performed KEGG pathway enrichment analysis. We found that the genes are involved in a variety of metabolic pathways, including vitamin B6 metabolism, ferric enterobactin transports, biosynthesis of siderophore group non-ribosomal peptides, cysteine and methionine metabolism, pantothenate and CoA biosynthesis, and some unclassified pathways (Table 1). Out of these 29 genes, 5 of them are involved in *E. coli* vitamin B6 biosynthesis pathway (Table 1). In contrast to eukaryotes, *E. coli* is capable of synthesizing the active form of vitamin B6, pyridoxal 5^′^-phosphate (PLP), through both *de novo* and salvage biosynthesis pathways (Figure 5A). In order to investigate the role of these two pathways in regulating the growth of worms fed with *L. plantarum*, we fed the worms with various *E. coli* mutants and evaluated their growth by measuring worm size. Our results indicate that the development of worms arrested at early stage when fed with a trace amount of *E. coli* mutants with disrupted vitamin B6 *de novo* biosynthesis (Figure 5B). This suggests that *E. coli* vitamin B6 *de novo* biosynthesis pathway plays an important role in recovering early-stage larval growth in the context of *L. plantarum* diet. To eliminate the possibility of contamination of these *E. coli* mutant strains, we performed PCR using gene-specific primers and verified the correct gene deletion in the knockout mutants (Figure S1).

**Figure 5.**
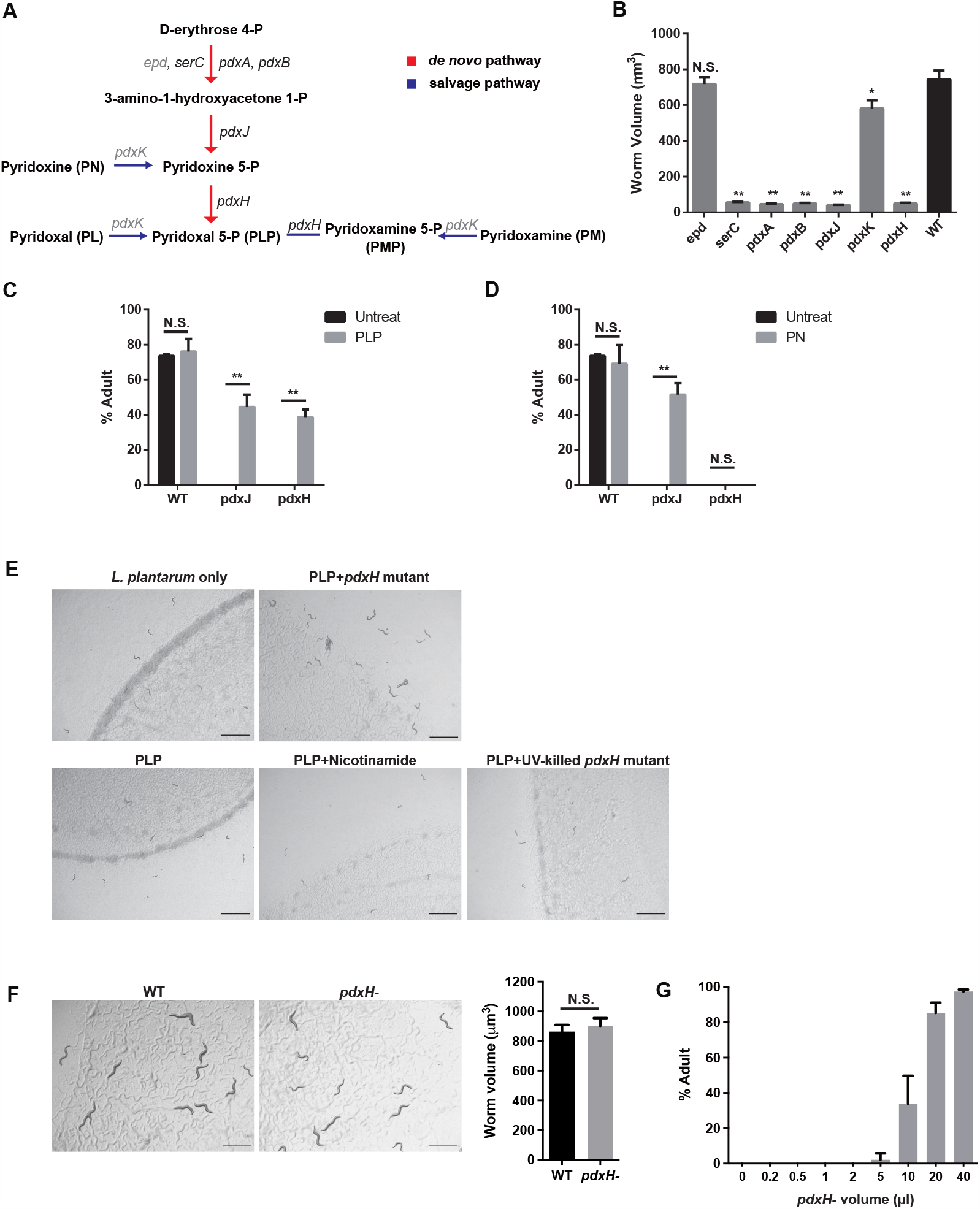

To further investigate the *E. coli* metabolites involved in vitamin B6 metabolism regulating arrested larvae development, the intermediate pyridoxine (PN) and final metabolite pyridoxal 5^′^-phosphate (PLP) were supplemented to the worms fed on trace *E. coli pdxJ* or *E. coli pdxH* mutants together with plenty of *L. plantarum*. We found the worm development completely recovered by dietary supplementation of the *pdxH* downstream metabolite PLP (Figure 5C, Figure S2). Meanwhile, supplementing with intermediate PN rescued worm growth on *pdxJ* mutant but not *pdxH* mutant (Figure 5D, Figure S2), indicating that the final product PLP is essential for *C. elegans* larval development. To determine whether the growth effect is restricted to a specific bacterial background, we added PLP only, PLP supplemented with the cofactor Nicotinamide, and PLP with trace amounts of UV-killed *E. coli pdxH* mutant to the worms that were fed on *L. plantarum*, individually. We found that supplementing 1mM PLP without live *pdxH* mutant did not promote the worm growth (Figure 5E), suggesting that the beneficial role of PLP for larval growth is likely dependent on some downstream bacterial usage of PLP. As trace amounts of *E. coli pdxH* mutant cannot support the growth of worms fed on *L. plantarum*, we investigated whether a plentiful amount of the *pdxH* mutant alone could support worm development. We placed worms on an *E. coli pdxH* mutant lawn and used BW25113 lawn as the control. The results show that the lack of *pdxH* does not affect worm growth (Figure 5F). Next, we explored whether other components from *E. coli* could compensate for the function of PLP and support the growth of worms fed on *L. plantarum*. To investigate this, we supplemented worms with a *L. plantarum* diet with varying concentrations of the *pdxH* mutant. We found that a supplementation of no less than 5 µl of *pdxH* mutant (OD600=1) could partially recover the growth of the worms (Figure 5G). This suggests that the function of *pdxH* can be compensated for by increasing the amount of the *E. coli* mutant. Taken together, these results demonstrate that PLP plays a role in recovering arrested larval growth with nutrient deficiency, and that the function of PLP can be partially compensated for by increasing the amount of *E. coli pdxH* mutant.

### PLP acts as the cofactor of host tyrosine aminotransferase and promotes L1 growth

To investigate the signaling mechanism that leads to developmental arrest of *C. elegans* fed on *L. plantarum* due to the absence of *pdxH* in *E. coli*, we carried out RNA sequencing and expression analysis to identify the host pathways regulated by *E. coli pdxH* activity. Specifically, we compared the RNA-seq results of worms fed a *L. plantarum* diet with either a trace amount of *E. coli pdxH* mutant or wild type *E. coli* BW25113. We identified 120 genes with increased expression and 830 genes with reduced expression with a Gfold (0.01) value (which indicates the fold change in gene expression) over 0.5 (S1 Dataset), suggesting widespread effects of the lack of *pdxH* in *E. coli* diet on host gene expression in *C. elegans*. Gene ontology (GO) analysis showed that upregulated genes were mainly involved in the metabolic process and oxidation-reduction process (Figure 6A), while downregulated genes were mainly involved in development and growth (Figure 6B), consistent with the developmental arrest phenotype. To gain insight into the crosstalk of vitamin B6 biosynthesis in *E. coli* with host vitamin B6 metabolism, we assessed the expression level of genes involved in host vitamin B6 metabolism. The RNA-seq results showed fluctuated gene transcription in host PLP binding (Figure 6C). Further qPCR analysis confirmed the relative mRNA level changes of two representative genes *eppl-1* and *F26H9*.*5*, compared to the internal reference gene *act-1* (Figure 6D). These results suggest a fundamental role of bacterial vitamin B6 in regulating host PLP binding activity and L1/L2 development.

**Figure 6.**
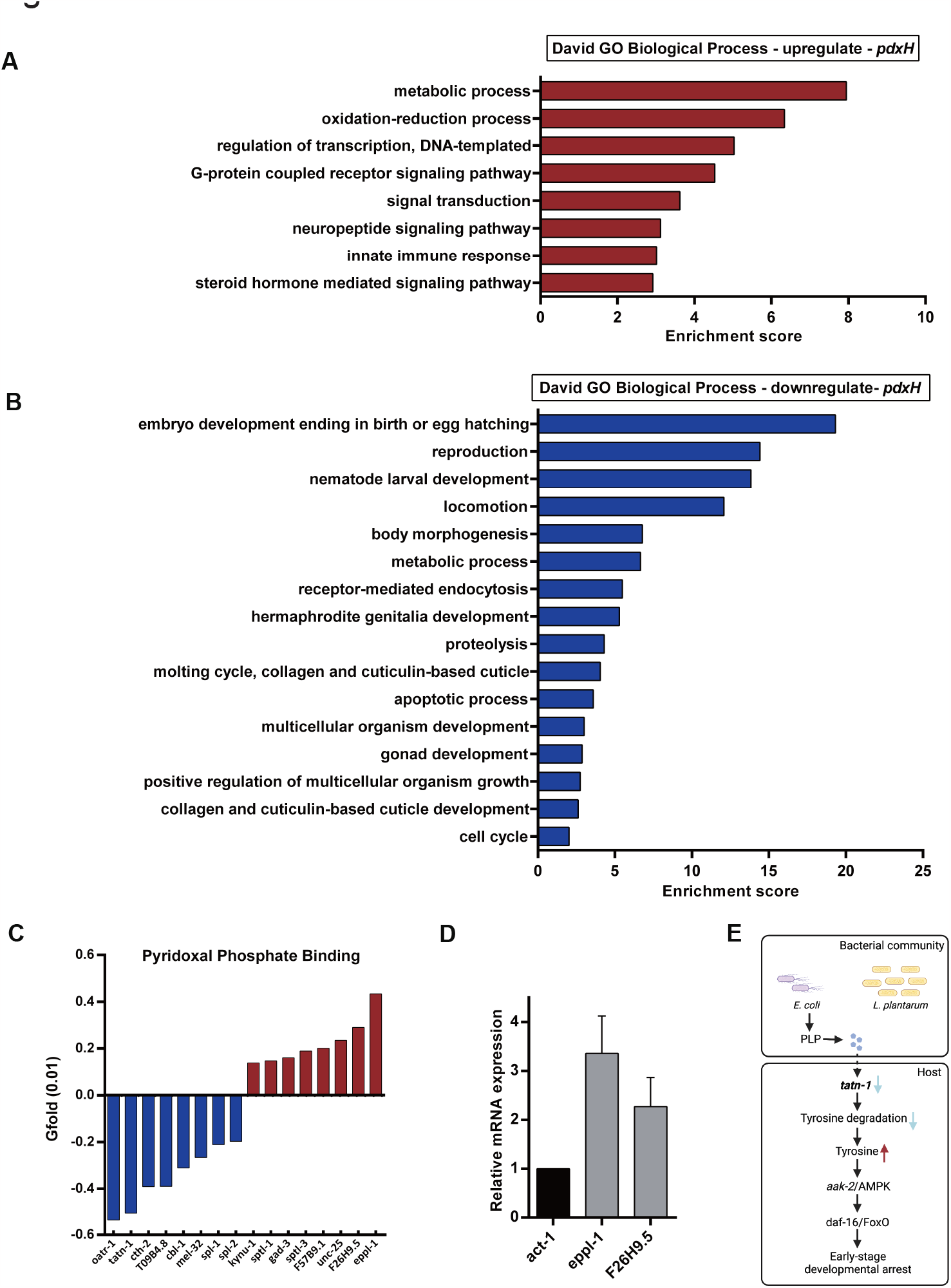

Interestingly, others have previously observed that the decreased expression of *tatn-1*, a gene encoding a tyrosine aminotransferase (TAT), results in reduced removal of tyrosine through degradation, leading to an increase in free tyrosine levels in *C. elegans*^12^. The free tyrosine then activates aak-2/AMPK and has a positive effect on daf-16/FOXO, resulting in the translocation of DAF-16 to the nucleus. Our RNA-seq analysis revealed a downregulation of the *tatn-1* gene in developmental arrested worms grown on *L. plantarum* diet supplemented with *pdxH E. coli* mutant. Previous studies have shown that TAT is a PLP-dependent enzyme that initiates the catabolism of tyrosine^13^. We hypothesize that the PLP from *E. coli* functions as a coenzyme of worm TAT, and the absence of PLP from the bacterial community leads to a reduced expression of tatn-1. The reduced clearance of tyrosine by TATN-1 leads to the accumulation of tyrosine, which activates aak-2/AMPK, causing the translocation of DAF-16 to the nucleus, and subsequently leading to the developmental arrest (Figure 6E). Overall, our findings suggest that bacterial PLP likely acts as a cofactor of host TATN-1, promoting worm growth through attenuating DAF-16 translocation.

## Discussions

Animals require essential nutrients to support their development. Microbes, either through commensal interactions or as food sources, provide certain essential nutrients that cannot be synthesized by the animal. Our study shows that *L. plantarum*, when provided as a nutrient-deficient food source for *C. elegans*, can induce reversible developmental arrest. This finding enhances our understanding that larvae hatching in the presence of food but lacking essential nutrients fail to initiate postembryonic development. Notably, a trace amount of *E. coli* enables worms to overcome this developmental arrest and complements the nutrient deficiency in *L. plantarum*. With this *L. plantarum*-induced developmental arrest model, we screened an *E. coli* single deletion library and found that the alteration of bacterial genetic composition influences *C. elegans* recovery from developmental arrest. As a result, the bacterial *pdxH* is not nutritionally adequate for the development of worm fed on *L. plantarum*. However, the supplementation of the *pdxH* downstream metabolite PLP compensates the nutrient deficiency and recovers developmental arrested worm growth. These results emphasized the importance of coexisting *E. coli* acts synergistically with *L. plantarum* in the biosynthesis of PLP and provides essential nutrients to initiate early-stage larval development in *C. elegans*.

In most animals, including *C elegans*, the PLP biosynthetic pathway is dysfunctional, requiring them to obtain vitamin B6 from their diet or microbiome^14^. Colonization of bacteria that has genomic diversity in vitamin B6 synthesis in *C. elegans* is beneficial to the health and life traits of the host^15^. In a chemically defined medium, *C. elegans* maintenance medium (CeMM), the active form of bacterial vitamin B6, PLP, is indispensable on worm population growth and health^16^. Regarding significance in development, bacterial PLP is essential to initiate early-stage larval development in the context of a *L. plantarum* diet (Figure 5D), however it is not required for the growth of further developed stages of worms, such as dauer worms (Figure 2F).

The ability of organisms to sense their nutritional environment enables them to alter their growth and metabolism accordingly. Previous studies on larval arrest have shown that the insulin/insulin-like growth factor signaling (IIS) pathway plays a crucial role in sensing the nutritional environment and regulating entry into arrest^17^. The IIS pathway is the principal regulator linking nutrient levels to metabolism and development in *C. elegans*. DAF-2, which is an ortholog of the insulin/IGF-1 transmembrane receptor, plays a key role in regulating L1 worm development^18^. DAF-16 is a widely expressed transcription factor belonging to the Fork head family, it regulates the expression of various genes involved in development, longevity, and stress resistance^18^. DAF-2 functions primarily through the regulation of DAF-16 to control worm developmental arrest^19^. As we gain a deeper understanding of the regulatory network controlling post-embryonic development, we may discover additional signals and signaling centers in the future. Given that *Lactobacillus* is also a commensal bacterium in many other metazoans, including humans, investigating the interaction between *Lactobacillus* and *C. elegans* may offer new insights into evolutionarily conserved probiotic effects.

## Methods

### Nematode and bacterial strains

Wild type N2 Bristol was obtained from the Caenorhabditis Genetics Center (CGC) and maintained on nematode growth medium (NGM) 6-cm Petri dish plates at room temperature. All experiments were performed with synchronized L1 stage hermaphrodite animals. *E. coli* OP50 and the keio collection parent strain *E. coli* BW25113 were grown at 37□ in Lysogeny broth (LB) Miller’s version. *E. coli* deletion strains were grown at 37□ in LB with 50 µg/ml kanamycin. *Lactobacillus plantarum* ATCC8014 was grown at 37□ in Lactobacillus MRS broth for 24 hours. Bacterial cultures were seeded on NGM plates and dried for 1 hour at room temperature prior to use in experiments.

### CFSE staining

The green fluorescent dye 5(6)-Carboxyfluorescein diacetate succinimidyl ester (CFSE, MedChemExpress) was used to track the bacteria. 1 ml of *L. plantarum* cultured 24 hours was harvested and centrifuged at 4000g for 5 min. Then resulting pellet was resuspended in 1 ml of 10 µM CFSE in PBS and incubated in the shaker at 37□ at 200 rpm for 30 min while avoiding light. After staining, the bacteria were washed three times with PBS and resuspended in PBS.

### Dauer larvae isolation

The N2 hermaphrodites were grown with OP50 at room temperature for approximately 10 days, during which there was an abundance of dauer larvae on the plate. The worms were then washed off the plate, and the worm pellet was resuspended in 1% SDS (sodium dodecyl sulfate). The worms were incubated in 1% SDS for 30 min with gentle agitation. Then, the worms were washed 1-5 times with M9 buffer to remove all SDS. To remove the carcasses of non-dauer larvae, 6ml of 30% ice-cold sucrose was added to the worm pellet and then centrifuged at 4□ for 5 min at a speed of 3,000 rpm. The floating worms were aspirated, and the pellet was washed twice with M9 buffer to remove excess sucrose. To eliminate the potential existing OP50 in the gut, the isolated dauer worms were washed twice with 100 µg/ml gentamycin, each wash lasting 10 min. After washing three times with M9 to remove gentamycin, the worms were suspended in M9 buffer.

### *E. coli* Keio collection screen

The MRS broth-cultured *L. plantarum* was concentrated to OD600=10 and resuspended in M9 buffer. Subsequently, 150 µl of *L. plantarum* suspension (1.2×10^8^ c.f.u.) was added to one side of peptone-free NGM plate. The *E. coli* mutants were cultured overnight at 37□ in LB medium with 10 mg/ml kanamycin. Then, 0.2 µl of each mutant culture (OD600=1, 1.6×10^5^ c.f.u.) was seeded to the other side of *L. plantarum* plate. Approximately 300 synchronized L1 worms were added to the screen plate and placed at room temperature. Then worm sizes were scored at day 3 and 4. Each mutant in the library was screened once for primary screen. For the secondary screening, 421 candidate mutants were screened in triplicate to confirm the slow growth phenotype.

### PCR verification of deletions

Single gene deletion *E. coli* strains were streaked onto LB agar plates containing 50 µg/ml kanamycin. Subsequently, PCR was performed on individual colonies using genomic and kanamycin-cassette-specific primers. Genomic primers were designed specifically for each strain at a distance of 100-300 bases upstream of the start codon of the gene, and 100–300 bases downstream of the stop codon. The primers used are listed in Table S1. The PCR products were then analyzed for the correct size by agarose gel electrophoresis.

### Metabolites supplementation

Pyridoxine (PN, Fisher BioReagents, 1mM) and pyridoxal 5^′^-phosphate (PLP, Thermo Scientific, 1mM) were dissolved in water and added to *E. coli* mutants and then spotted on to the plates.

### Total RNA extraction

*C. elegans* at the same stage grown on *L. plantarum* together with trace amount (0.2 µl, 1.6×10^5^ c.f.u.) of wild type BW25113 or *pdxH* mutant were collected from peptone-free NGM plates and washed three times with M9 buffer. Then 250 µl of TRIzol reagent (Invitrogen) was add to the worm pellet and worms were homogenized for 5 min. RNA was isolated by adding 50 µl of chloroform, followed by centrifugation to separate the aqueous phase. The aqueous phase was transferred to a new tube and mixed with isopropanol (125 µl) to precipitate RNA. The resulting RNA pellet was washed twice with 70% ethanol (250 µl). RNA pellets were air-dried and then re-suspended in RNase-free water. Any potential genomic DNA contamination was removed by DNaseI treatment using the TURBO DNA-free Kit (Invitrogen).

### Reverse transcription quantitative PCR (RT-qPCR)

RT-qPCR was performed using the CFX Opus 96 Real-Time PCR System (Bio-rad). To begin, total RNA samples were reverse transcripted into first-strand cDNA using the iScript Select cDNA Synthesis kit (Bio-rad). 1 µl of a 10X diluted cDNA sample was then used in the qPCR reaction with SsoAdvanced Universal SYBR Green Supermix (Bio-rad). Primers were designed using Primer3 software (v4.1.0). Relative mRNA expression levels were calculated using the 2^-ΔΔct^ method, and the reference gene act-1 was used to normalize the gene expression data.

### RNA-seq

Purified RNA was quantified and assessed for quality using Epoch 2 microplate spectrophotometer (BioTek). cDNA libraries were prepared from 500ng RNA per sample. Libraries were then sequenced on an Illumina HiSeq 2000 sequencing machine using paired-end sequencing with a read length of 100 nucleotides. Adaptor sequences and low-quality reads were removed using Trimmomatic, The RNA-Seq reads were then mapped to the *C. elegans* genome using STAR 2.5.3a with default settings. Transcript abundance, measured as read counts per gene, was extracted using HTSeq. Differential gene expression analysis was performed using GFold^20^. Gene ontology (GO) analysis was conducted using DAVID.

## Statistical analysis

The statistical analyses were conducted using GraphPad Prism and Excel. The results are presented as the mean ± SEM, and the data were evaluated using an unpaired two-tailed Student’s t-test.

## Declaration of interests

The authors declare no competing interests.

## Acknowledgments

This work was supported by the Texas A&M Startup grant from TEES and Department of Chemical Engineering to Q.S.

## Notes

https://data.mendeley.com/drafts/cmvv6y66dz

### Competing Interest Statement

The authors have declared no competing interest.

