## Supplementary figures and images for "Bacterial vitamin B6 required for post-embryonic development in *C. elegans*"

### Supplemental Figures

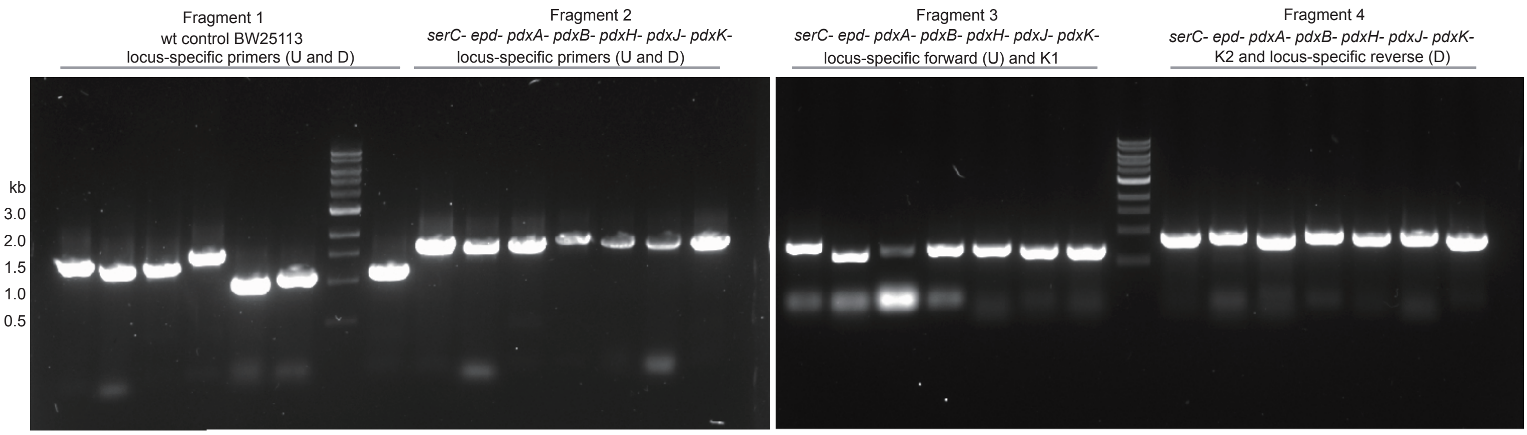

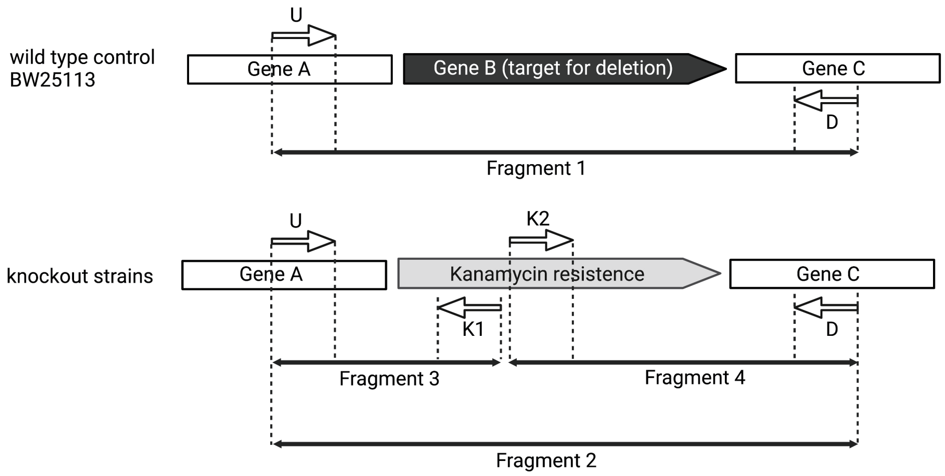

#### Figure S2

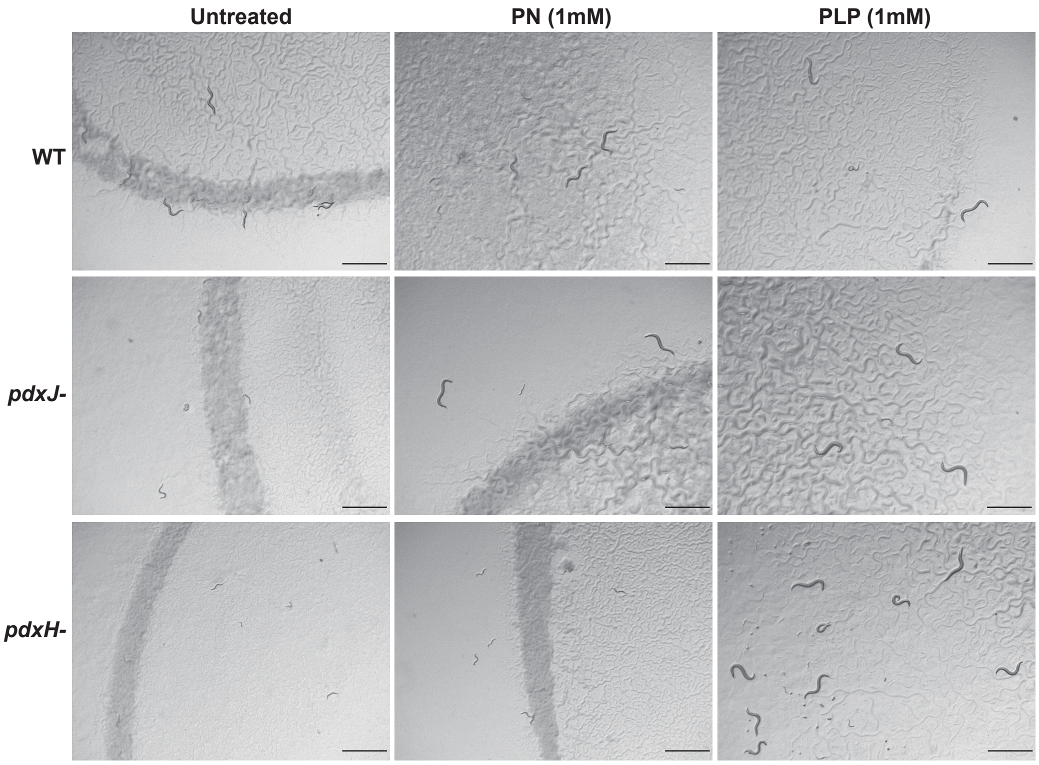
